# Widespread analytical pitfalls in empirical coexistence studies and a checklist for improving their statistical robustness

**DOI:** 10.1101/2023.07.04.547661

**Authors:** J. Christopher D. Terry, David W. Armitage

## Abstract

1. Modern Coexistence Theory (MCT) offers a conceptually straightforward approach for connecting empirical observations with an elegant theoretical framework, gaining popularity rapidly over the past decade. However, beneath this surface-level simplicity lie various assumptions and subjective choices made during data analysis. These can lead researchers to draw qualitatively different conclusions from the same set of experiments. As the predictions of MCT studies are often treated as outcomes, and many readers and reviewers may not be familiar with the framework’s assumptions, there is a particular risk of “researcher degrees of freedom” inflating the confidence in results, thereby affecting reproducibility and predictive power.
2. To tackle these concerns, we introduce a checklist consisting of statistical best-practices to promote more robust empirical applications of MCT. Our recommendations are organised into four categories: presentation and sharing of raw data, testing model assumptions and fits, managing uncertainty associated with model coefficients, and incorporating this uncertainty into coexistence predictions.
3. We surveyed empirical MCT studies published over the past 15 years and discovered a high degree of variation in the level of statistical rigour and adherence to best practices. We present case studies to illustrate the dependence of results on seemingly innocuous choices among competition model structure and error distributions, which in some cases reversed the predicted coexistence outcomes. These results demonstrate how different analytical approaches can profoundly alter the interpretation of experimental results, underscoring the importance of carefully considering and thoroughly justifying each step taken in the analysis pathway.
4. Our checklist serves as a resource for authors and reviewers alike, providing guidance to strengthen the empirical foundation of empirical coexistence analyses. As the field of empirical MCT shifts from a descriptive, trailblazing phase to a stage of consolidation, we emphasise the need for caution when building upon the findings of earlier studies. To ensure that progress made in the field of ecological coexistence is based on robust and reliable evidence, it is crucial to subject our predictions, conclusions, and generalizability to a more rigorous assessment than is currently the trend.

## Introduction

The quest to understand how similar species can avoid competitive exclusion is a central goal of community ecology (Chase & Leibold, 2003; Hutchinson, 1959) has generated an extensive body of theory describing the formal conditions required for pairwise coexistence (Amarasekare, 2019; Barabás et al., 2018; Chesson, 2000). A key strength of this approach, frequently referred to as Modern Coexistence Theory (MCT), is to unite the niche and neutral theories into a consistent theoretical framework (Adler et al., 2007). It defines coexistence as a balance between niche differences and relative fitness differences with the mechanisms giving rise to these differences named *stabilising* and *equalising* mechanisms, respectively. In simple theoretical models, such as the Lotka-Volterra competition model, these quantities can be derived using simple ratios of parameters. The combined theoretical simplicity and conceptual power of this approach to unify our understanding of competition and community assembly (Grainger, Levine, et al., 2019) led to its widespread and accelerating adoption as a framework around which to design experiments investigating the roles of species interactions in community assembly.

The standard approach to empirical MCT comprises a series of steps that are tractable for individual research groups to undertake: using small-scale experiments or natural population dynamics in defined plots to parameterise density-dependent population growth functions. The model’s parameters are then used to calculate niche and fitness differences which predict a binary coexistence outcome based on a simple, theoretical inequality. Importantly, experiments can be repeated under different controlled treatments to identify shifts in these key quantities and subsequent coexistence predictions. This general approach has been followed by dozens of studies and is expanding rapidly. Such studies have been conducted across a wide range of taxonomic groups (e.g. Mediterranean annuals (Matías et al., 2018), insects (Terry et al., 2021), yeasts (Grainger, Letten, et al., 2019) and pondweeds (Hess et al., 2022)) as well as experimental treatments (e.g. rainfall (Van Dyke et al., 2022), temperature (Armitage & Jones, 2020), consumers (Petry et al., 2018) and shared pollination (Johnson et al., 2022)).

At its core, the empirical utility of the MCT framework hinges on its assumed ability to correctly classify, through prediction, the categorical outcome of coexistence or exclusion. Predictions and analysis of coexistence are typically made through satisfaction of the well-known inequality defining the potential for mutual invasibility: *ρ* < *κ*_1_/*κ*_2_ < *ρ*^−1^ where *ρ* is niche overlap and *κ*_1_/*κ*_2_ is the relative fitness differences between the two competitors. These approaches require an assumed functional form of inter- and intraspecific density dependence (such as the Lotka-Volterra or Beverton-Holt models) and data from a competitive arena — which can be experimental or observational — between two species. This fitted model is used to predict each competitor’s maximum intrinsic growth rate and competition coefficients. Other taxon-or model-specific values may also be collected, such as seed germination fraction or dormancy rates.

However, empirically characterising competition, even between pairs of species, remains a considerable practical challenge (Hart et al., 2018; Inouye, 2001). Despite the apparent simplicity of the results described in the MCT framework, there is no best single, benchmarked data analysis pipeline for its empirical applications. Instead the process of data analysis involves a multitude of decisions that are both data- and model-dependent, best characterised as a ‘garden of forking paths’ (Gelman & Loken, 2014). This flexibility encompasses a wide range of choices, such as selecting from comparable candidate models or determining thresholds for defining ‘significant’ effects. As a result, the influence of researcher degrees of freedom in shaping conclusions may be as substantial as the data itself. Coexistence inferences are based on indices of model parameters that are both individually hard to empirically estimate and frequently confounded leading to the potential for bias propagation. Despite frequently following the same fundamental recipe, there has been a wide divergence in the level of statistical rigour to assess these issues across studies (Fig 2), resulting in diminished support for the hypotheses under investigation (Armitage, 2022; Terry, 2023).

**Figure 1.**
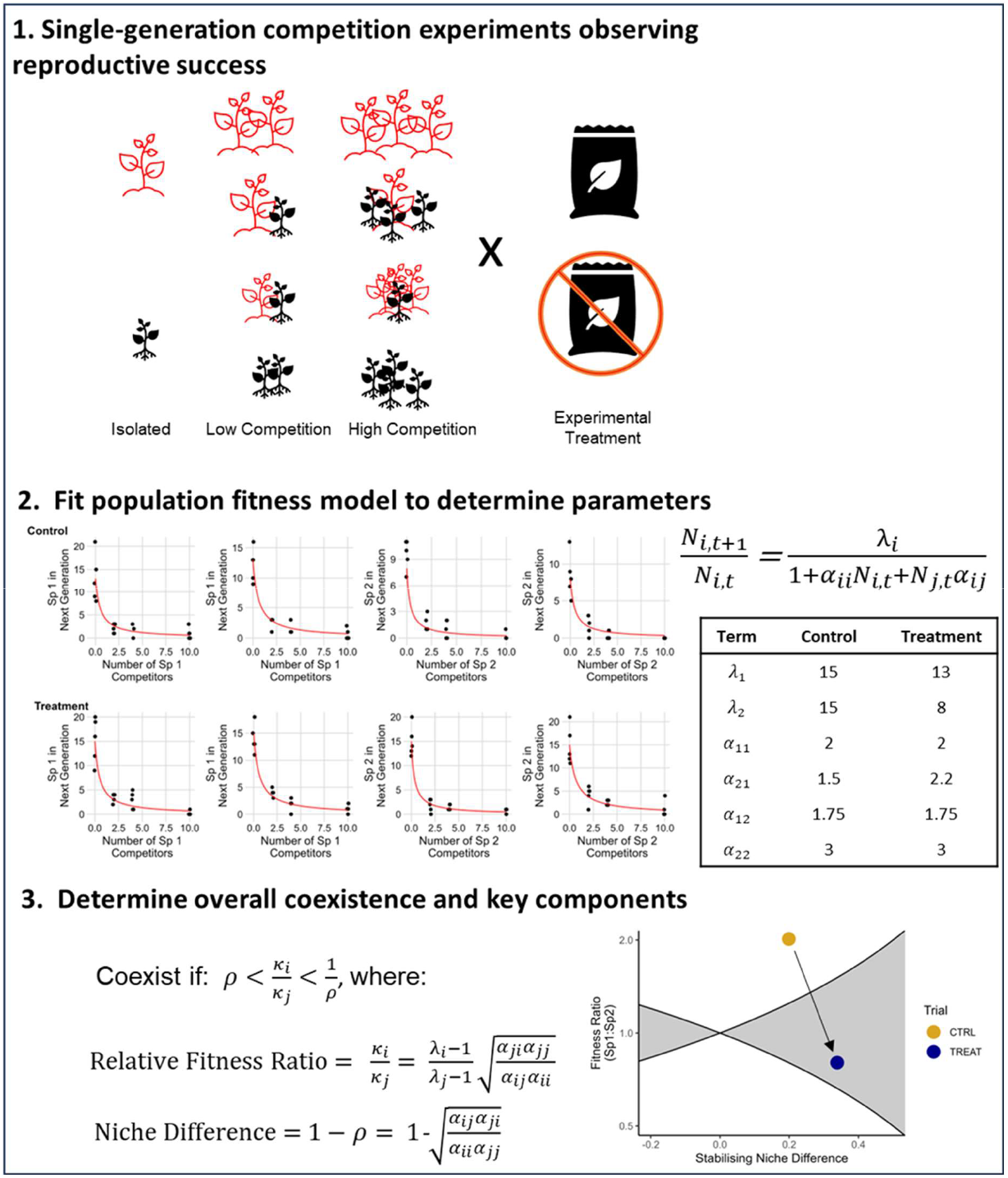
Illustration of the three steps common to most empirical MCT studies. Here, researchers ask whether a treatment affected the predicted coexistence between a pair of species. In Step 1, a large number of population fitness assays of both species are run under different levels of competition and levels of the treatment. In Step 2, a model is fit to this data (here a Beverton-Holt model) to predict the per capita population growth rate per generation. Note that for this idealised example, the data was simulated from this model with Poisson error but the plotted fit curves in red are generated using the ‘true’ values of the parameters listed in the table. Lastly, in Step 3, predicted coexistence or exclusion is assigned using a theoretically-motivated inequality. These identify the conditions for coexistence and label particular quantities as relative fitness differences (RFD) and stabilising niche differences (ND) which can be summarised in the coexistence plane plot. In this case, the treatment is identified as shifting the system from the exclusion of species 2 to predicted coexistence.

**Figure 2.**
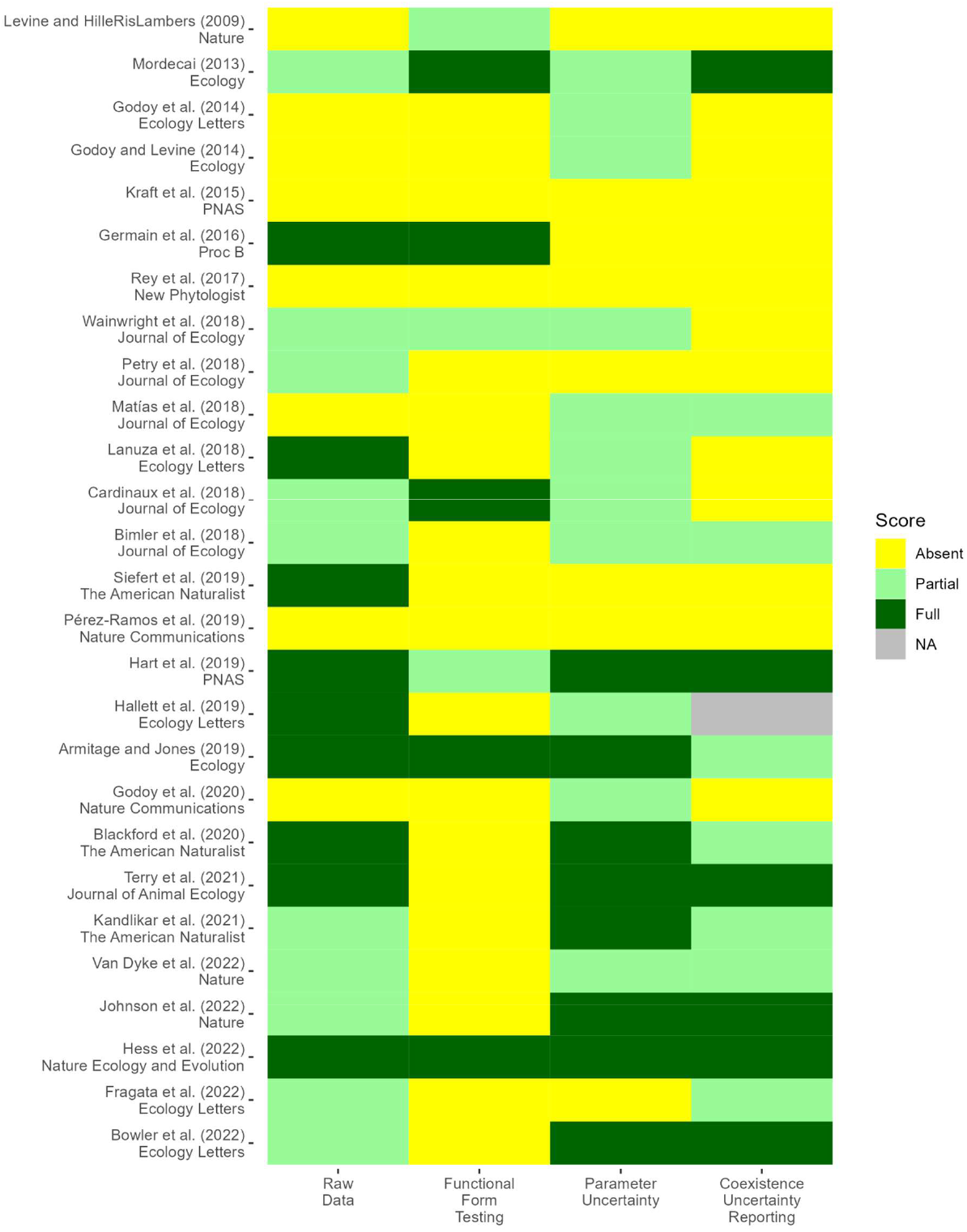
Summary of select studies that make use of the competitive response surface approach described in Figure 1. A more detailed table, including studies using different approaches such as invasion analysis or fitting to time-series data is included in the accompanying repository. Assessment of whether studies partially or fully meet criteria is necessarily somewhat subjective. The four columns align to the four categories in our checklist: a) Raw data: is the raw observed data available and is the model fit plotted with the raw data? b) Functional form testing: Do the authors test alternative competition models and fully report the model comparison? c) Parameter uncertainty: Do the authors report the uncertainty in the competition model parameters? Studies reporting just standard errors were graded as ‘partial’ and full confidence intervals as complete. d) Coexistence uncertainty reporting: is the final coexistence outcome reported or tested as a single point estimate, are error bars of some sort reported (partial), or is uncertainty propagated fully to the end result?

Here we present a short checklist of recommended statistical steps and an overview of these recommendations as they relate to making predictions in empirical coexistence studies. While checklists may be perceived as inflexible to authors’ preferred experimental and analytical approaches, this is a mischaracterisation of their intended use. A checklist is not a prescription, but rather a prompt for active consideration of one’s analytical steps and a reference point for reviewers less familiar with the methods (T. H. Parker et al., 2018). While each check is intended to be uncontroversial and consistent with standard protocols for presenting models fit to empirical data (Zuur & Ieno, 2016), they are frequently omitted in published papers. We therefore highlight with examples how seemingly innocuous omissions can lead to erroneous or unsupported conclusions specifically within empirical MCT. The experimental work required to parametrise the models is demanding, but this should increase the motivation for statistical rigour to make best use of hard-won data. Handling multiple forms of uncertainty simultaneously represents a significant challenge across scientific fields (Milner-Gulland & Shea, 2017; Simmonds et al., 2022), but strong tools exist that can significantly enhance the robustness of results.

All ecological theory requires a set of starting assumptions in order to be operationalised and deployed (Grainger et al., 2021). However, successful deployment of theory in an empirical setting requires these assumptions to be regularly tested. For MCT these assumptions can be split into two categories. First, that competition between the focal species is well described by one’s chosen model and the parameters can accurately generate reliable inferences. Second, that the validity of the theoretical framework itself holds in real-world scenarios. These latter assumptions may include the suitability of a pairwise model in multispecies settings (Barabás et al., 2016; Levine et al., 2017), the negligible role of demographic stochasticity in coexistence assessment (Pande et al., 2020; Schreiber et al., 2023), and the lack of interference from confounding environmental factors. Here we focus on this first set of assumptions concerning model suitability as they are directly relevant to ongoing trends in the empirical MCT literature and are often a point of confusion by readers unfamiliar with the MCT approach.

There are a range of alternative approaches to applying MCT in empirical settings (Godwin et al., 2020)and differing interpretations of niche and fitness differences (Spaak et al., 2023). For consistency, our discussion and examples cover the widely-used experimental design and interpretations outlined in Figure 1. However, many of the conclusions also apply under alternative definitions of niche and fitness differences (Carroll et al., 2011; Narwani et al., 2013) or approaches that fit models to long-term time series data (Adler et al., 2006).

### BOX 1.

Checklist Summary

1. Availability and accessibility of underlying data
  a. Raw data and computer code are permanently, publicly available in a standard format
  b. Raw observations are plotted alongside fitted model predictions
2. Selecting the core population growth model
  a. Basis for selecting the functional form(s) of density-dependent responses is reported
  b. Choice of data transformations and error distributions are justified
  c. Comparison and selection among alternative candidate model(s) has been carried out in line with study objectives
3. Addressing uncertainty in model coefficients
  a. Uncertainty around parameter estimates is reported
  b. Support for the assumption of treatment effects on parameters is presented
4. Propagation of full uncertainty to final results
  a. Parameter uncertainty is propagated into predicted coexistence metrics (niche and fitness differences, invasion growth rates)
  b. Coexistence predictions are interpreted in light of this uncertainty

BOX 1. Summary of key checklist items for robust empirical inferences within the Modern Coexistence Theory framework.

## 1. Raw experimental data

### Check 1a - Availability of raw data accompanied by metadata

The recent shift towards publishing raw observational data underlying experimental results has been a major advance for reproducible science (Culina et al., 2018; Reichman et al., 2011). Publication of raw original data (typically in comma or tab-delimited text format), alongside reasonably annotated code and metadata, has key advantages over just publishing tables of fitted parameter values. Sharing raw data not only enables the complete replication of the original methods, but also facilitates the implementation of novel analytical approaches and the testing of emerging theories (Jenkins et al., 2023). Fortunately, an improvement in raw data availability is a clear trend in our survey of empirical MCT studies (Figure 2), although we encountered recent examples where datasets were described as open, but not linked to from the paper. Relatedly, sharing of computer code, ideally annotated, greatly aids method transparency and reproducibility.

### Check 1b - Raw data plotted with best fit lines

Despite being a requirement in introductory statistics classes, the graphical presentation of raw data alongside modelled trend-lines is surprisingly uncommon (Figure 2). This is unfortunate, as such plots are useful for assessing the accuracy and precision of nonlinear population models. The human eye is excellent at pattern detection, and visual assessment underlies the more quantitative checks we describe in the following sections. Even where the raw data is archived it should not be expected that the average reader or reviewers generate these plots themselves. While it can be awkward to present raw data from large experiments, in most cases these plots can be moved to supplementary material to avoid space restrictions.

## 2. Model-dependence of results

There are a variety of mathematical functions describing how an individual’s reproductive output is affected by various forms of competition. Choosing among many similar, alternative competition models is a longstanding problem in ecology (Ayala et al., 1973; Law & Watkinson, 1987; Martyn et al., 2021), but is a requirement of most approaches for predicting coexistence (Cervantes-Loreto et al., 2023), since all coexistence metrics and their associated uncertainties are contingent on the model itself being a reasonable representation of reality. Each population’s monoculture equilibrium and their invasion growth rates when the competitor is at that equilibrium are quantities central to MCT’s coexistence metrics. Most often, rather than being directly observed, they are derived from extrapolations of a fitted model. Because of this, relatively small changes in functional form or parameter values can result in qualitatively different predictions about coexistence. It is therefore surprising how few of the surveyed studies explicitly compare or present fit statistics of the models being used to make inferences.

Competition models used in MCT studies tend to be phenomenological (that is, the myriad mechanisms contributing to pairwise competitive outcomes are collapsed into a single coefficient), and are often nested or very similar in parameters and complexity. Because of this, we argue that there is usually no *a priori* “best” model for a given system, and even if there is, this choice should still be justified by comparing its fit statistics to plausible alternatives. Model comparison is an active area of statistical research and though it can be challenging to offer universal advice, in general, we suggest the very act of comparing multiple candidate models is a critical step, regardless of the specific metric or approach taken. In this section we will consider how two aspects of the model fit to raw data– the functional form of the response to competition and the error structure can influence results and offer some guidance about the process of model selection in an MCT context.

### Check 2a – Justification of the competition model’s functional form

Across our literature survey, models were frequently described as having been chosen based on simplicity and use in prior studies. In particular, studies of coexistence in annual plant communities nearly unanimously assume the Beverton-Holt model best describes the dynamics of the systems (SI table). This simple model assumes a saturating form of density dependence (Figure 1) from which it is convenient to derive niche and fitness differences. Specifically in the context of MCT — which relies on extrapolating fitted models to equilibrial abundances — reasonable alternative model structures can lead to drastically different inferences of predicted coexistence (Abrams, 2022; Cervantes-Loreto et al., 2023). An example of this using simulated data is shown in Figure 3.

**Figure 3.**
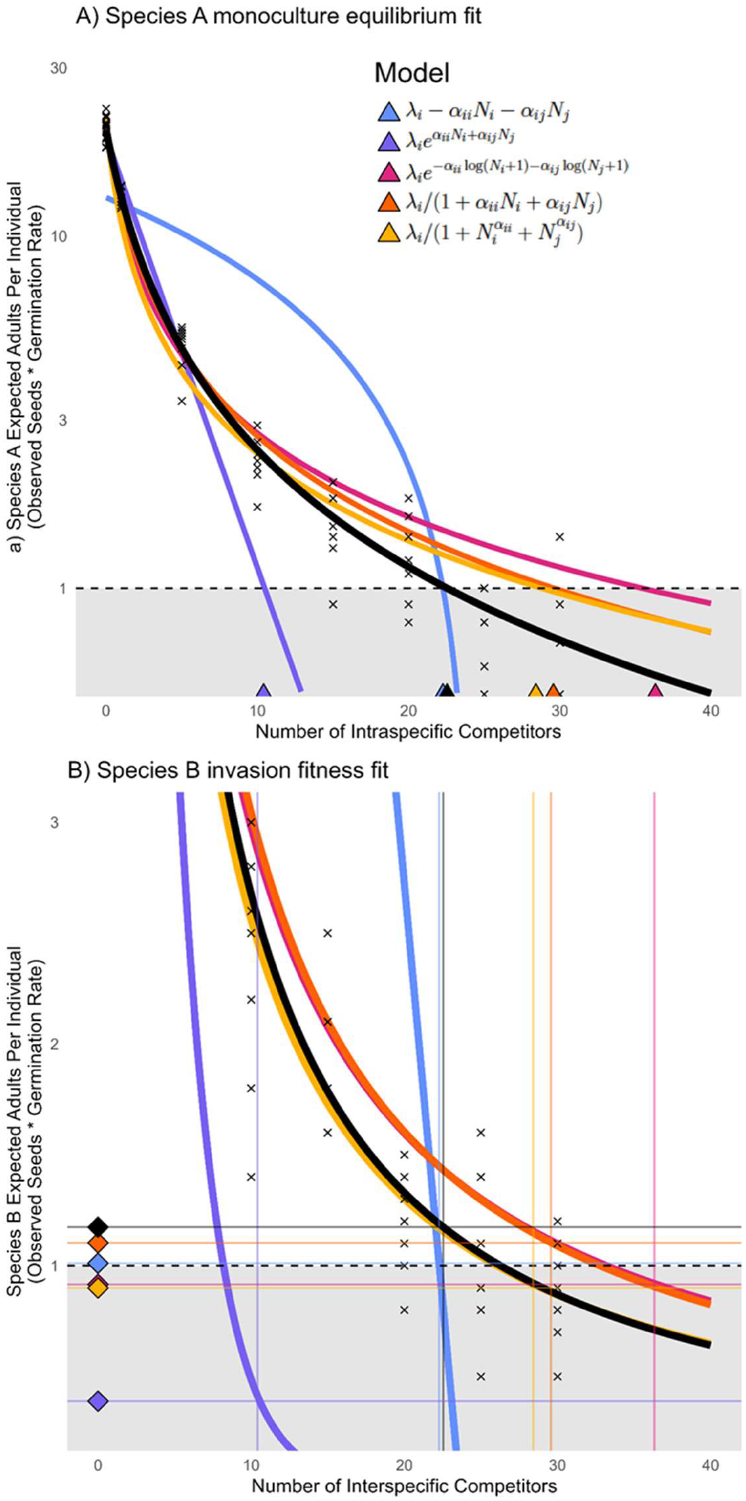
Illustration of influence of model specification on coexistence predictions. Here, simulated data is generated using a model where the reduction in seed production is specified with a three-parameter expression 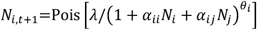. See code supplement for parameter values. Five standard competition models are fit using *brms* to a moderate amount of simulated data. Poisson noise on the observed seed-counts was included, but note that the overall amount of noise in these simulated data is considerably less than most real examples. Data and the ‘true’ model are shown in black. A) Data and model fits to estimate the monoculture equilibrium of Species A 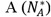, as calculated by the number of intra-specific competitors that give an expectation of 1 replacement individual in the next generation (black horizontal line). Visually, three models appear to fit well, but give markedly higher estimates of 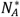. Note that y-axis has been logged to highlight differences. B) Data and mean model predictions to estimate the expected population growth rate of Species B at the predicted monoculture equilibrium of A. Where above 1, B is said to be able to invade A (and so potentially coexist if the reverse is also true). Note that the y-axis has been zoomed to highlight this key quantity. In this example only Model 5 is able to identify that species B should be able to invade species A and, in this case, model comparison by ELPD was able to identify Model 5 as the best fit.

When competition trials are not held at densities at or beyond competitors’ carrying capacities, there is an additional imperative to select a competition model that can reasonably be extrapolated to estimate the zero-net-growth conditions at which invasion growth rates are calculated. In such situations direct model selection may have limited capacity to correctly differentiate models that give very different predicted equilibrium values and it would be necessary to select a model based on precedent and acknowledge the dependence of the results on these choices.

A key first step in this process is defining the set of models to be compared. Since most competition models are phenomenological, there is essentially no limit to the forms that could be tested, and there are strong arguments that more mechanistic generative models should be developed for specific well-studied groups of organisms (e.g., annual plants, Stouffer, 2022). However, when phenomenological models are all that is available, they tend to fall within the Volterra-Lotka-Gause family of models, which enjoy reasonably straightforward derivations of niche and fitness differences from their shared parameters (e.g. those listed by Law & Watkinson, 1987). Because of this, identification of alternative candidate models should also consider flexibility in the shape of density-dependent responses (see (Novak & Stouffer (2021a) for a discussion of issues in the similar context of fitting consumer functional response models). We suggest that the diversity of functional forms is more important than the total number of candidate models. Further comparison of competition models with null, competition-free or single-coefficient competition models can also provide helpful context on the appropriateness and fit of the corresponding fully parameterised model. When the raw data follow a different shape than that stipulated by candidate models it may be necessary to develop new derivations that better match reality (Godoy & Levine, 2014).

### Check 2b - Justification of error structure and transformations

Experimental or observational data always contain error components that fitted competition models can only imperfectly capture – the exact same combinations of competitors often result in a wide range of fecundities and competitive effects among identical replicates. This error is frequently modelled as observation error, though in most cases, the measurements (e.g. counts of seeds) are likely to be quite accurate. Instead, inter-replicate variation frequently results from process error in the sense that is caused by real ecological mechanisms such as intraspecific variation, microclimate differences and stochastic events which differentially affect experimental replicates. Furthermore, it is common for the focal species in competition experiments to have different reproductive strategies and hence different variability between species. Certain plants, for example, produce extremely variable numbers of seeds per individual and within a species the distribution of seed production can also vary from normally-distributed to highly-skewed or zero-inflated. This variation (and how it is handled) has consequences both for observed population dynamics (Hart et al., 2016; Pascual & Kareiva, 1996) and the fitting and assessment of competition models.

In this context, careful consideration needs to be given to data transformations, error distributions, and whether the variance is treated separately for each species. The core problem is that the mean of many transformations of a random variable is not generally equal to the transformation applied to the mean -i.e. Jensen’s inequality. There has been much discussion of this issue in other ecological contexts (Richards, 2008; Warton et al., 2016), yet we found it barely mentioned in our survey of the MCT literature. Because the goal of the model fitting stage is to precisely and accurately approximate parameter values, rather than to simply assess their statistical significance, generic advice and assumptions concerning transformations from a direct null-hypothesis testing context may be less relevant.

Until somewhat recently, least-squares and maximum likelihood (ML) approaches to parameter estimation were necessitated by computational constraints. In the specific context of nonlinear competition models and noisy measurements, these methods can be unstable and result in convergence errors unless transformations are first applied to the data. In particular, fitting Gaussian models to log-transformed seed counts has been a common approach in plant competition models (Godoy et al., 2014; Law & Watkinson, 1987) to normalise the variance and make non-linear models easier to fit. This can improve model likelihoods when data is sparse, but introduces biases for MCT beyond the widely appreciated challenge of zero-values. The most meaningful ‘average’ value of seed count from a single generation is an arithmetic mean, not the geometric mean estimated from a fit to the logarithm of seed counts. As such, where the variation is ‘real’ (i.e. not just measurement error) fitting to log(seeds) will underestimate the expected mean number of seeds per individual of that species by a factor proportional to *exp*(*σ*^2^).

Particularly where the variance differs between species, these biases in estimates of maximum growth rates (*λ*) and competition (*α*) can have significant consequences for predicted coexistence outcomes. In the simulated example presented in Figure 4 where data are lognormally distributed, the fitted parameter values to the logged data markedly underestimated the population growth rate terms and overestimated the competition terms, particularly of species 1, which had the higher error. This seemingly minor decision to transform the data has the consequence of completely switching the predicted competitive hierarchy from the “true” outcome. For an example using real data, see Supplementary Figure 1. In that example, both a Poisson and a Negative Binomial error distribution were able to give good predictions, but the Poisson model was overly confident in its parameter estimates.

**Figure 4.**
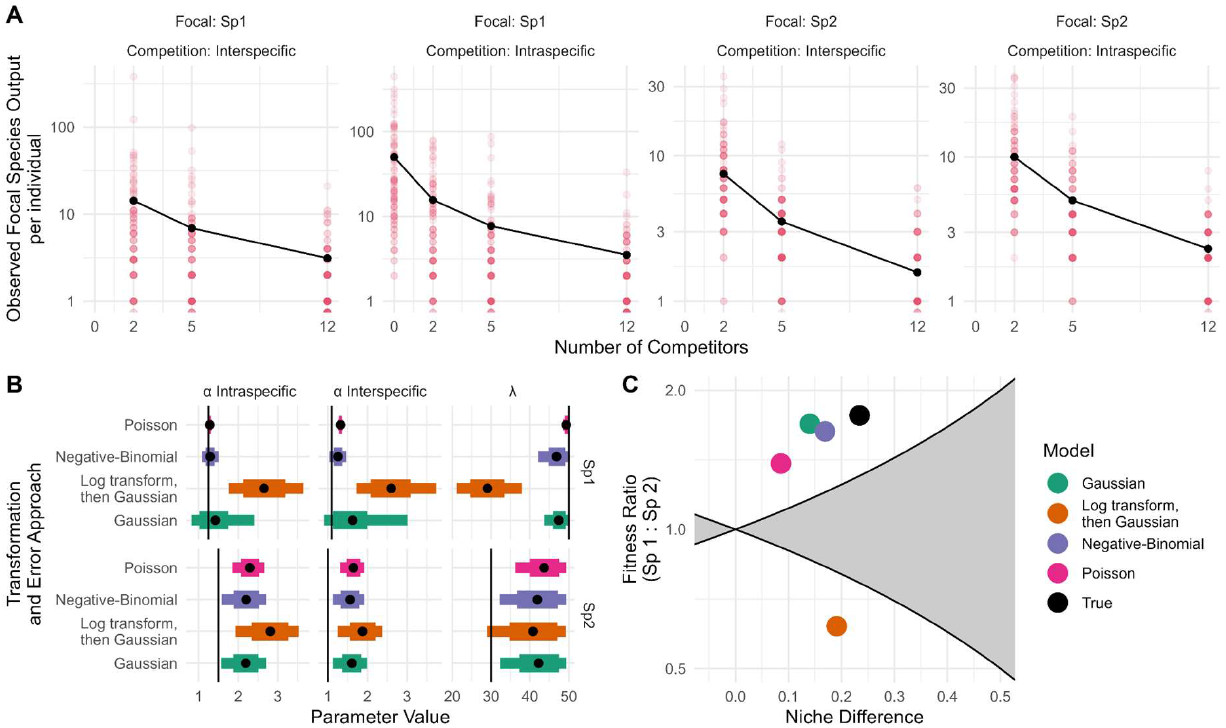
Demonstration of impact of data transformation and error-structure on coexistence predictions. Here observation data of two species (1 and 2) is simulated from a Beverton-Holt model (parameters *λ*_1_= 50, *λ*_2_ = 30, *α*_11_= 1.1,*α*_22_= 1, *α*_21_= 1.5, *α*_12_= 1.25). A) Observations (100 per competitor level) were drawn from a discretized log-normal distribution (*log σ*_*sp* 1_= 1.3, *log σ*_*sp* 2_= 0.6) with an expected mean value equal to that predicted by the competition model (black lines). B) Four different error approaches were used to fit the ‘correct’ Beverton-Holt competition model using *brms* to these data: i) directly fitting a Gaussian error model to raw observations, ii) log-transforming (log(x+1)) the observations then assuming Gaussian errors, iii) fitting a Poisson error model to raw observations, iv) fitting a negative-binomial error model to raw observations (fitting separate shape parameters for each species). Parameter estimates are shown with means, central 95% and 60% intervals. The ‘true’ values used to generate the data are shown with the vertical black line. C) Coexistence plane diagram showing predictions of the four models, and the ‘true’ location.

The development of efficient Monte Carlo approaches for model fitting (for example STAN (Stan Development Team., 2022) and its accessible R frontend *brms* (Bürkner, 2018)) mean there is more flexibility in directly modelling error and avoiding the need for transformation. Error models can account for the functional form of uncertainty such as the commonly-observed heteroskedasticity in offspring production across gradients of competitor densities. Error models can also be fully pooled (i.e. a single error term fit) or separated for each focal species and treatment level. For counts, a negative binomial or a quasi-Poisson error structure can capture more complex and non-symmetric error patterns, while estimating mean effects on their natural scales.

That said, it is rarely easy to choose an error model *a priori* solely on theoretical and biological grounds. Visual inspection of model fit, particularly by plotting the distribution of residual errors, can often be helpful to determine candidate models for residual variation. If comparison of different functional forms is undertaken, it is little additional work to include alternative error distributions for each candidate model. Care must be taken to compare the data on the same response scales, since many statistical programs automatically use non-identity link functions when certain families of error distributions are specified. In these cases, each model’s parameters must be back-transformed before coexistence metrics can be calculated.

### Check 2c Model comparison and selection

It is well established, although often taken for granted, that the goals and outcomes of model selection depend on the objectives of the study (Aho et al., 2014; Tredennick et al., 2021). This is mostly widely understood in terms of the classic metrics: Akaike’s information criterion (AIC) is generally preferred for its capacity to identify a model with the best predictive power, and the Bayesian information criterion (BIC), which penalises free parameters more than AIC, is favoured for accurately identifying the “true” model. Although the implementation differs in a Bayesian framework, for example using LOOIC (Leave-One-Out-Information Criteria, Vehtari et al. (2017)) or WAIC/WBIC (Watanabe-Akaike/Bayesian information criterion Watanabe (2013)), fundamentally the same issues of selection for pure prediction versus inference remain. Although model comparison and selection can be accomplished in a variety of ways, we recommend all include a reporting of the range of models examined, the criteria used to identify the best supported models and (where there is not a clear model) robustness of the conclusions to the use of reasonable alternative models.

For a typical experimental coexistence study (Figure 1), the overall predictive ability of the underlying fecundity model is likely to be less relevant for most questions than accurately describing the functional form of competition, which could imply a BIC-family measure would be preferred.

Within a classical maximum likelihood framework, BIC is as easy to implement as AIC. However, at the time of writing for Bayesian models there is considerably better support for model selection on the basis of predictive performance (such as WAIC and ELPD), in particular through the ‘loo’ R package (Vehtari et al., 2022) for STAN models. Another more direct approach to model fitting uses cross-validation either among experimental replicates or generations to identify the best competition model. However, because most coexistence studies suffer from low replication and are rarely conducted over more than one generation, cross-validation is rarely performed. Despite this abundance of somewhat arbitrary methods to choose from, engaging closely with and reporting the *process* of model selection is more important than the specific methods used.

Where unique, non-nested models are equally supported, repeating the full analysis using multiple model structures can be informative. If the end conclusions are consistent, then this is strong additional support for the conclusions (e.g. Hess et al. (2022)). By contrast, if the conclusions are contingent on a particular model, when alternative functions also fit the data well (Armitage, 2022), then additional data (or justification) may be required to support one over the others. When dealing with nested models, numerous models will share equivalent parameters and can be used on a species-specific basis, since their derived coexistence metrics have the same formulae. More flexible forms of niche and fitness difference metrics also exist (Spaak & De Laender, 2020) that can directly compare invasion growth rates estimated from non-nested models, sidestepping the need for the same model or even the same model family to be assumed for all species.

Formulaic use of model selection criteria is not without its own risks. Model selection algorithms cannot replace the visual assessment of different models across and beyond the range of data. In particular, inspection of the pattern of residuals across the data is helpful to identify cases where predictions could be dependent on the sampling design of the raw data. Where simple models are favoured by information criteria, consideration should be given to the likelihood that this is a symptom of noisy data, rather than the generality of parsimonious models. In particular, inference based on parameter estimation after model selection can be a problem (Yates et al., 2023) as estimates of significance can be positively biased. Model selection issues have been identified as causing systematic bias in the analysis of predation functional response models which share many of the same fundamental difficulties as competition models (Novak & Stouffer, 2021b).

Whichever model selection path is followed, for clarity and to catch errors it is helpful to explicitly report both the fitted log-likelihood (or its equivalent) and number of fitted parameters for each model. More heavily parameterised models are prone to identifiability issues and can struggle to converge with noisy data (Hess et al., 2022; Levine & HilleRisLambers, 2009). For truly nested models, increased model complexity should only increase the log-likelihood. Hence for each parameter removed, it would be expected that the maximum improvement in AIC is 2. Results that do not follow this pattern strongly suggest that the model fitting has not been successful.

## 3. Parameter Uncertainty

Point estimates of the parameters of competition models are never exact owing to the statistical uncertainty derived from measurement error and real intraspecific variation due to differences in micro-environment, genetic, epigenetic and parental effects. This intraspecific variation can in itself have significant consequences for species coexistence (Hart et al., 2016; Stump et al., 2022).

Theoretical studies have investigated the sensitivity of coexistence to perturbations to key parameters (Barabás et al., 2014). However these sensitivity analyses are applicable to mathematically small perturbations – i.e. those that don’t shift populations into alternative dynamical regimes such as from persistence to extinction.

Beyond the mechanistic effects of variation in growth parameters, the model fitting process itself returns variation around point estimates of model parameters. Appraisal of this variation is a critical but often-missing aspect of many coexistence experiments. In practice, the number of competitive trials per free parameter is frequently moderate (typically ranging from 10-20 across studies, although sample sizes were not always reported). Nonetheless, some degree of residual uncertainty is expected given the limitations of time- and labour-intensive experiments – for many ecological questions it is often advantageous to test more species pairs reasonably well rather than exhaustively characterise competition between a single pair. The question is how to appropriately evaluate and work with this residual uncertainty to generate reliable and actionable predictions of coexistence.

### Check 3a – Report Uncertainty in Model Coefficients

Our literature review revealed that a majority of studies included some reporting of the uncertainty in the estimates of key competition model parameters— most frequently as standard errors. This is helpful to allow both readers and reviewers to assess the confidence affordable to the results, but there is still room for improvement. The likelihood profiles of competition (*α*) terms are often non-symmetric, and are poorly summarised by a single value. Where feasible, it is advantageous to either provide 95% intervals or ideally plots of the posteriors. However, an important caveat is that the challenges of the separability of parameters in competition models causes the uncertainty associated with a particular parameter to be dependent on others. In particular, *λ*_*i*_ and *α*_*ij*_ estimates are frequently correlated, with consequences for MCT predictions that are not easily summarised in a list of parameter values (see next section). Further, standard confidence intervals for parameters implicitly rely on the underlying model’s suitability, and so can give artificially high confidence when alternative models have not been compared. For instance, in the example in Figure 4, the Poisson-error model gave very precise but inaccurate estimates for the competition coefficients.

### Check 3b Report support for Effect of Treatments on Model Coefficients

An objective of many MCT studies is to identify the effect of an experimental treatment on coexistence through its actions on the metrics of niche and fitness differences. In most cases, these derived metrics are calculated from ratios of demographic parameters fit from the competition models. Some go further, for example Van Dyke et al. (2022) compare the effect of a water treatment on the two component parts of the relative fitness differences: the demographic ratio 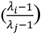 and the competitive response ratio 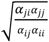. However, it is important to assess whether the experimental data statistically support the assumption of treatment effects on parameters in the first place, to avoid conclusions being unduly influenced by noisy data.

A key challenge is that interaction coefficients between species (*α* terms) are more difficult to accurately estimate than intrinsic reproductive rates (*λ*). This is usually caused by fewer numbers of data points being available to estimate the competition parameters. Experimenters can easily include relatively large numbers of isolated individuals to directly assay *λ* terms whereas *α* terms rely on multi-species experiments across a gradient of relative frequencies. In designs with more than two species, every trial with the focal species can contribute to the estimation of *λ*, while only those between specific pairs can assess *α* terms. Further, forcing treatment effects in one’s model fitting processes can introduce bias and uncertainty, as doing so further reduces the number of data points available to estimate each parameter. Consequently, the influence of individual, noisy data points are more likely to influence *α* terms compared to *λ* terms, particularly when separate parameters are being fit for each experimental treatment (Terry, 2023). In light of this, how should a practitioner assess whether the assumption of treatment differences in model parameters are warranted?

One approach is to compare models that fit separate parameters between treatments with models where the parameter values are shared between treatments. However, in larger experiments with more than a single pair of species, the appropriate scale at which to test treatment effects can be hard to define, particularly for the competition coefficients. For instance, it is not always clear if assessment of treatment effects should be conducted for each *α* separately (as per Johnson et al. (2022)), for all *α*’s belonging to each focal species (Terry, 2023), or over all terms (e.g., both *α*’s and *λ*’s) simultaneously (as per Terry et al. (2021))?. Comparing models in this manner can highlight cases where individual species experience treatment-driven shifts in growth or competition, but practitioners must be cautious of the increasing false positive risk with increased numbers of species. Ultimately, the approach taken and choices made concerning whether to accept or reject specific treatment effects hinges on a pluralistic assessment of statistical evidence for or against these effects combined with expert system-based judgement. If, for example, treatment effects show no increase in a model’s predictive performance but large confidence margins, then one might be better served by using all of the available data to fit a single competition parameter with higher confidence.

Alternatively, simulation analysis can identify the sensitivity of the analytic pipeline and end results to noise in the empirical data. Where appropriate, simulated data that does not include a treatment effect can be used as a null model against which to test the effect of hypothesised shifts (Terry, 2023). This approach is particularly useful in translating uncertainty across levels from population-growth model parameters through to final results, where direct analytical solutions to uncertainty propagation are challenging.

## 4. Handling and Propagation of Uncertainty

Errors around parameter estimates do not disappear after being used to calculate coexistence metrics. Rather, propagation of this error through to the final coexistence predictions is critical for a fair assessment of the evidence in favour or against coexistence. While error around individual model parameters is frequently reported, only a small number of papers fully propagate this uncertainty into their coexistence predictions. Handling model uncertainty is a significant challenge across scientific fields (Simmonds et al., 2022). In an MCT context, error propagation into final coexistence predictions has been used to inform the results (Cervantes-Loreto et al., 2023; Hart et al., 2019; Hess et al., 2022; Mordecai, 2013; Terry et al., 2021). Bowler et al., (2022) demonstrates particularly well how a failure to propagate uncertainty can result in misleading conclusions about coexistence.

When fitting competition models, the parameters are often not fully separable (i.e., they are non-identifiable), resulting in highly correlated estimates of growth and competition parameters. Since most of the key quantities of MCT are ratios of parameters, these covariances can influence the overall error. While propagating error is best achieved by using Bayesian posterior draws of the parameters to fit distributions of coexistence metrics, it can also be done in a frequentist framework through methods such as moment approximations via Taylor expansions or Monte Carlo simulation (Dietze, 2017). In most cases, the additional work is relatively straightforward. For example, when using the R language, propagating error is often as simple as plugging in a randomly-sampled set of posterior draws for each parameter in place of a single value. In the frequentist case, when only a point estimate and associated standard error is returned, error is most easily propagated into niche and fitness difference equations using packages such as *propagate* (Spiess, 2018)-the principal challenge is often in reporting the results in concise and readable format.

As a further incentive, propagating multivariate uncertainty when parameters covary can often tighten the confidence in the final predictions. Where variables are divided, positive covariance decreases overall variance. Hence, the most likely covariance arising between parameters in competition experiments — between *λ*_*i*_ and *α*_*ii*_ or *α*_*ij*_ — will tend to decrease uncertainty in the relative fitness ratio comprising the fitness differences coexistence metric.

### Check 4a – Propagation of uncertainty to coexistence plane graphs

The central results of many coexistence studies are presented on cartesian coordinates depicting the predicted outcome of competition between a pair of species in terms of the niche and fitness differences. Here, zones of ‘coexistence’ and ‘exclusion’ are demarcated based on the coexistence inequality discussed in the introduction. Plotting outcomes on this plane provides an accessible visual summary of the predicted consequences of experimental treatments on coexistence. However, there are currently no conventions for the appropriate visualisation and interpretation of error on such graphs.

Figure 5 illustrates the range of approaches that have been used to present results on a coexistence plane. While two orthogonal error bars around niche and fitness metrics (Fig 5b) are preferable to nothing, they have multiple deficiencies hindering their interpretation. First, the true distribution of uncertainty is often very irregularly-shaped. This arises from covariances between underlying parameters as well as the fact that the *α* terms contribute to both coexistence metrics, making the two axes non-independent. Secondly, interpretation of this 2D error requires evaluating the extent to which uncertainty is distributed across a nonlinear boundary defining the predicted outcomes, which can be difficult to assess with only two orthogonal error bars (Figure 5b). Full posterior distributions of coexistence metrics display the most information on uncertainty (Fig 5d), but can be overwhelming when many species pairs or treatments are compared on the same plane (Fig 5c). An underused alternative is to use small pie charts (Yan et al., 2022) summarising the relative number of outcomes of coexistence, exclusion, or priority effects predicted by the posterior draws (Fig5e). Importantly, in most cases these plots can only be expected to convey a visual summary of the result, and statistical tests and summary statistics should be carried out and discussed. In many cases, there are considerable advantages and very little cost to including the posterior draws as a supplementary .csv file so that future users of the data do not need to refit the models.

**Figure 5.**
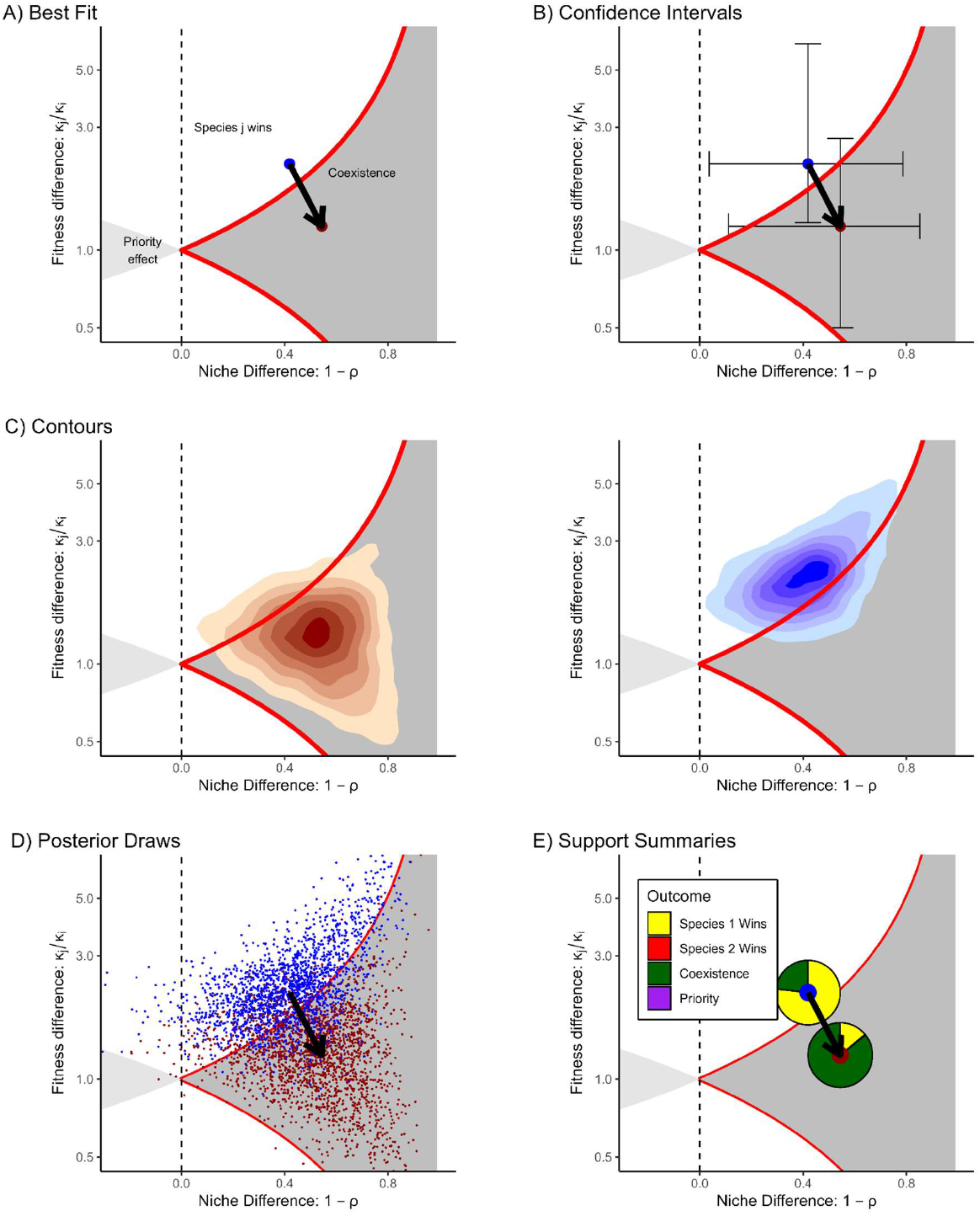
Options for representing uncertainty on coexistence diagrams. A) Plotting point estimates do not permit assessment of the statistical support for or against coexistence predictions. B) Orthogonal error bars (here 95% intervals) indicate a high uncertainty in predicted outcome but do not account for the underlying covariance between niche overlap and fitness differences. C) Graphical summaries of the posterior distribution, such as contour plots, can be more digestible than raw posterior draws but remain hard to quantitatively interpret and usually require facetting of subplots, increasing the space required. D) Plotting posterior draws directly display uncertainty while permitting comparisons between treatments but are less statistically interpretable than other options. E) Support summaries offer one way to both maintain the visual simplicity of changes in best fit while including the fraction of posterior support for each of the four potential predictions.

### Check 4b – Propagation of uncertainty to final overall results

We note with concern that highly uncertain niche and fitness differences have often been summarised as single, qualitative binary outcomes, even after the large uncertainties in their underlying parameters have been reported. Most papers presenting uncertainty do so with orthogonal error bars, but do not explicitly mention or interpret this uncertainty in their reported conclusions. In most cases this uncertainty is a formal expression of the uncertainty in the model, rather than a prediction for the frequency by which any given coexistence treatment or species pairing is likely to result in a particular categorical outcome. However, these interpretations are related — a pair of species with high intrinsic variability in pairwise outcomes will also be difficult to place in a binary coexist or exclude category.

Assessing the evidence for or against coexistence in an error-conscious manner allows the researcher, reviewers, and readers to fairly assess the degree of evidence for or against these qualitative, categorical predictions. Bowler et al. (2022) provide a clear application of this approach, wherein joint posterior distributions that have nontrivial densities on either side of the coexistence boundary cannot — under common definitions of statistical evidence — be reliably assigned to either category and must therefore be scored as an ambiguous outcome. While such a conclusion clearly detracts from the cleanliness of the results, we argue that as coexistence outcomes are being increasingly discussed in applied contexts such as restoration (Aoyama et al., 2022) and invasion biology (Epstein et al., 2019), presenting results without corresponding uncertainty is of limited value to managers or other decision-makers.

Beyond the issue of assigning coexistence outcomes in a single species pair or treatment, summarising the dynamic effect of experimental treatments on coexistence outcomes is challenging. While it is clear that illustrating treatment effects using only single summary points for each treatment can over-exaggerate these effects (Armitage, 2022; Terry, 2023), there is no statistical convention for assigning significance to treatment effects on predicted coexistence outcomes even when uncertainty has been quantified. To this end, rather than suggesting a hard-and-fast rule for identifying treatment-driven changes to coexistence (such as a <5% overlap in the two treatments’ joint posterior densities, for instance), we advocate practitioners to transparently justify their chosen reasoning for or against such shifts. Options to support such statements include Bayes factors, which quantify relative support for one hypothesis over another, given the data, without arbitrary significance thresholds.

While the presentation of probabilistic evidence can require large, ungainly tables (e.g. Terry et al., (2021)), a study’s main error-conscious conclusions can still be concise. For example, rather than stating that a treatment ‘*qualitatively altered predicted competitive outcomes for x of the n species pairs*’ based on the relative position of pairs of mean or median points, uncertainty can be directly incorporated into the conclusions with a rephrasing such as ‘*within our community, our results suggest for a random pair of species a p% probability that coexistence outcomes differed among treatments* ‘, or an alternative that most closely aligns with the study’s objectives.

In non-treatment designs, a continuous predictor such as trait or phylogenetic differences are used to predict niche or fitness differences (and hence coexistence, e.g. (Kraft et al., 2015; Pérez-Ramos et al., 2019). However, treatment of coexistence metrics in the standard regression framework ignores their underlying uncertainty and can artificially deflate true statistical effects. Instead, modern regression techniques such as error-in-variables models (Caroll et al., 2006) widely used in meta-analyses can incorporate previously-quantified uncertainty and more accurately identify statistical trends. Equally, approaches to quantifying coexistence mechanisms based on partitioning invasion growth rates (Ellner et al., 2019) can also include propagation of uncertainty in parameter estimates (Aoyama et al., 2022).

### Pre-study Considerations

While our proposed statistical checks all take place after experimental data have been gathered, we also recommend some steps be taken when planning experiments in order to avoid some of the aforementioned pitfalls. As a first step, we recommend pre-registration of coexistence experiments. Pre-registration entails drafting and archiving a report outlining the hypothesis, experimental plan, sample sizes, and analytical roadmap of one’s planned experiment (T. Parker et al., 2019). Such practices are becoming commonplace in many fields, and high-profile journals now offer provisional publishing agreements following adherence to peer-reviewed registered reports (Gya et al., 2023; “Rolling out Registered Reports,” 2023). At minimum, we advocate for practitioners to present more well-defined *a priori* hypotheses concerning treatment-mediated effects on coexistence. Rather than simply hypothesising that a treatment will affect coexistence outcomes in some unknown direction and magnitude, a stronger hypothesis would predict the direction of the effect and a clear criterion for how the effect is to be measured and supported.

A second design consideration is sample size since it influences the accuracy and precision of model parameters. Logistical and financial resources can constrain sample sizes, which risks exaggerating responses via type I errors, especially when combined with longstanding publication biases in favour of positive results (Yang et al., 2022). As the scope of most coexistence studies are multispecies assemblages, there is a tradeoff between the generality of the results and precisely estimating individual pairwise interactions. An assessment of the statistical power required to detect effects is a recommended approach to informing sample size requirements. To do so, one must already have an approximate guess of the target parameter values, in particular for the expectation of variability and effect sizes for treatments. When calculating power is not feasible, we suggest a minimum of at least 10-40 individual data points per parameter as a starting point for fitting competition models. For communities of moderate diversity (∼5-10 species) and two experimental treatments, this can require thousands of replicates to parameterise all pairwise interactions (Terry et al., 2021; Van Dyke et al., 2022). Efforts to reduce the dimensionality of the problem such that the number of experiments required no longer increases with the square of the number of species will be central to scaling-up empirical coexistence studies to the natural diversity a community (Stouffer et al., 2021; Weiss-Lehman et al., 2022).

## Conclusion and Outlook

Based on our literature survey, it appears that a significant number of published empirical MCT papers possess noticeable deficiencies in the statistical treatment of their data, which weakens the strength of their conclusions. Additionally, many foundational coexistence studies do not provide access to their raw data, hindering their re-analysis. Unfortunately, our case studies reveal that many seemingly innocent omissions, such as insufficient model comparisons or justifications, can relatively easily influence key ecological conclusions.

A recognised strength of MCT is the framework’s generalisability across a wide range of taxa. In principle, a similar experimental design could generate the data required to measure niche and fitness differences for any tractable group of organisms. This sets up the exciting potential for meta-analyses to investigate generalities in the relative importance of coexistence mechanisms (Buche et al., 2022; Yan et al., 2022). However, without individual studies reporting uncertainty in their coexistence predictions, their findings are not effectively usable by meta-analysts. Currently, it is difficult to move beyond the concern that the diverse range of conclusions regarding the relative importance of niche or fitness differences in facilitating coexistence may be influenced, at least partially, by researcher degrees of freedom.

Coexistence theory is on the precipice of moving from relatively abstract theory (Chesson, 2000), through pioneering initial demonstrations (Levine & HilleRisLambers, 2009) to hope for widespread adoption in empirical applications (Grainger, Levine, et al., 2019; Terry et al., 2022). In order to fully harness its potential benefits, it is essential that empirical studies stand on replicable and solid statistical foundations. Although this article offers many critical insights, we hold a strong sense of optimism regarding the future prospects of MCT in empirical research. We aim for this guide to contribute to charting a statistically rigorous course toward a productive and insightful future for the coexistence programme.

## Data Availability

This manuscript uses no new experimental data. All R code and data used to generate the figures is available at https://github.com/jcdterry/MCT_Review_public and archived on Zenodo at https://doi.org/10.5281/zenodo.8113527/

## Author Contributions

Both authors conceptualised the framework of the article. JCDT led the writing of the manuscript, initiated the literature review and made the examples. DWA contributed to the literature review and both authors contributed substantially to revisions and development of the manuscript.

---

**Figure S1.**
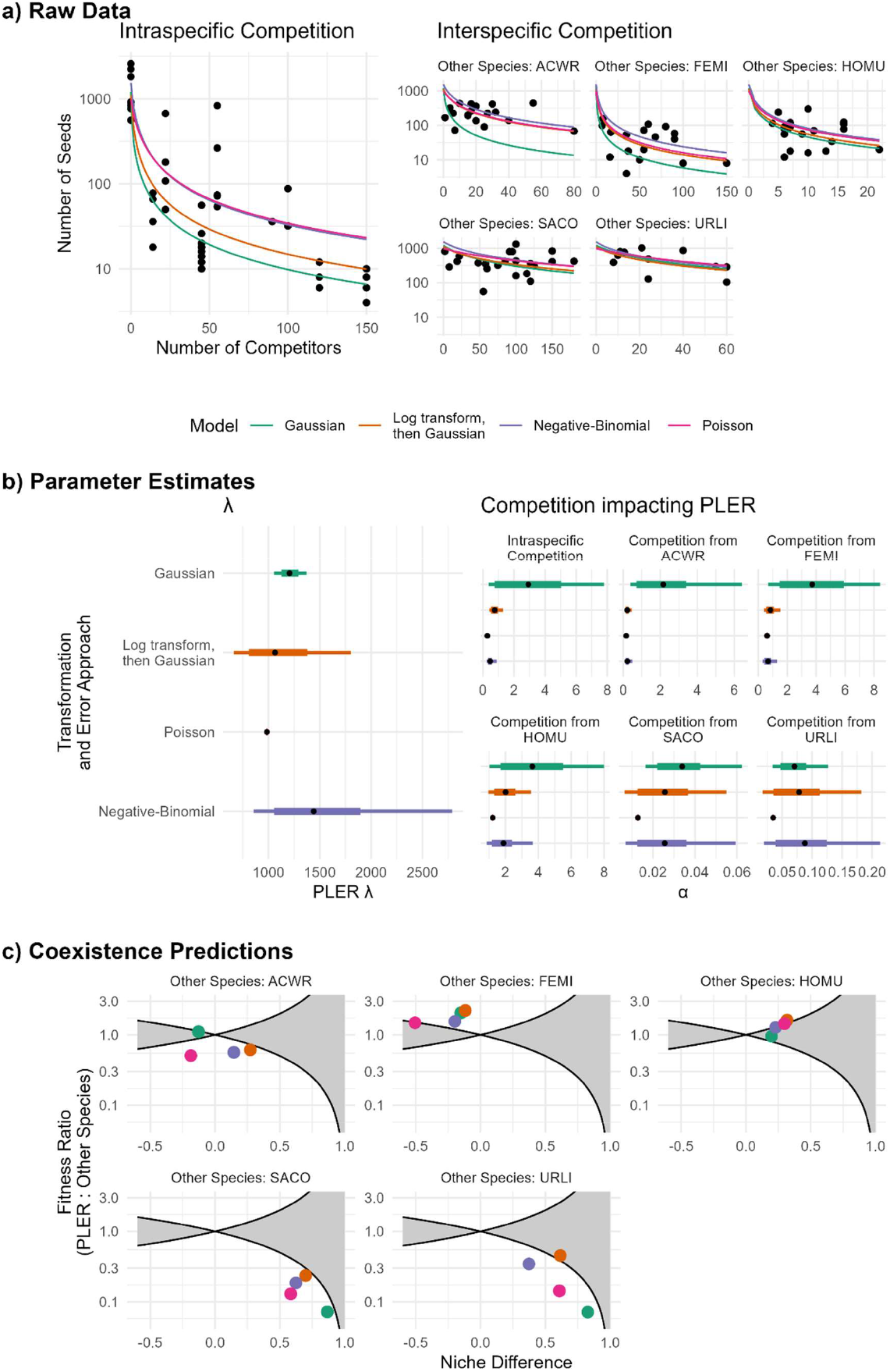
Impact of different approaches to modelling error on fitted parameters and predicted coexistence using real data. Data is sourced from Van Dyke et al (2022) for seed production of species *Plantago erecta* (‘PLER’) in the ‘wet’ treatment in competition with five other species coded ACWR, FEMI, HOMU, SACO and URLI. All approaches use the same Beverton-Holt competition model, but different error models or transformations. These models were Green: Directly fitting to the raw observed seed number with a Gaussian error model. Orange: Taking logarithms of both raw data and the prediction and then fitting with a Gaussian error model Pink: Fitting to raw data with a Poisson error model. Purple: Fitting to raw data with a Negative-Binomial error model. Different sigma or shape parameters were fit for each focal species. a) Raw data (black points) and fitted models (coloured lines). The raw data is highly skewed with occasional very large numbers of seeds produced – note the logarithmic y-axis. b) Similar to the simulated data used in the main text, estimated parameter values differed between the models in both their central estimate and their confidence. Parameter ranges show central 90% and 60% intervals and the black point the mean. Note that lambda was fit on a logarithmic scale c) Estimates of niche and fitness differences between PLER and the five other species, and hence coexistence predictions, change markedly between error models used. Models were fit in STAN via brms with loose priors -see code supplement for further detail.

